# Generalized Rayleigh-Plesset Theory for Cell Size Maintenance in Viruses and Bacteria

**DOI:** 10.1101/552778

**Authors:** Abdul N. Malmi-Kakkada, D. Thirumalai

## Abstract

The envelopes covering bacterial cytoplasm possess remarkable elastic properties. They are rigid enough to resist large pressures while being flexible enough to adapt to growth under environmental constraints. Similarly, the virus shells play an important role in their functions. However, the effects of mechanical properties of the outer shell in controlling and maintaining the sizes of bacteria or viruses are unknown. Here, we present a hydrodynamic “bubbles with shell” model, motivated by the study of bubble stability in fluids, to demonstrate that shell rigidity and turgor pressure control the sizes of bacteria and viruses. A dimensionless compliance parameter, expressed in terms of the elastic modulus of the shell, its thickness and the turgor pressure, determines membrane response to deformation and the size of the organisms. By analyzing the experiment data, we show that bacterial and viral sizes correlate with shell elasticity, which plays a critical role in regulating size.

## I. INTRODUCTION

Viruses consist of genetic material surrounded by a protective coat of proteins called capsids, which withstand high osmotic pressures and undergo modification (or maturation) to strengthen capsids after viral assembly (1). The protective coat is critical in enabling the virus to maintain its functionally intact state. In bacteria, the envelope covering the cytoplasm, besides being essential in sustaining the shape of the cell, protects the bacteria from adversary factors such as osmotic shock and mechanical stress (2–4). Bacterial cell wall is composed mostly of peptidoglycan, whose synthesis, regulation and remodeling are central to bacterial physiology (5). Growing body of evidence suggests that proteins controlling the organization of peptidoglycan growth could be crucial in the maintenance of cell size (2, 6). Bacteria and viruses exhibit remarkable diversity in size and shape. Nevertheless, for the purposes of developing a physical model, we picture them as spherical envelopes enclosing the material necessary for sustaining their lives.

Individual strains of bacteria are known to maintain a narrow distribution of size even when they divide multiple times (7–9). A number of physical models exploring how microorganisms maintain size and shape have been proposed (10–13), with similarities between cell elongation and bubble dynamics (10, 11). Historically, cell size maintenance has been discussed in terms of two major models: “timer,” where cells grow for a fixed amount of time before division, and “sizer,” where cells commit to division at a critical size (14). Another important model is the “adder” mechanism, which proposes that a constant size is added between birth and division (8, 15, 16). These models incorporate a ‘license to divide’ approach (17) - depending upon the passage of time, growth to a specific size or addition of fixed size to trigger cell division and regulate size. Recently, the need to attain a steady state surface area to volume ratio was proposed as the driving factor behind size homeostasis in bacteria (18). In emphasizing the need to move away from a ‘birth-centric’ picture, alternate models relating volume growth to DNA replication initiation have been proposed (9, 19) based on experiments (20, 21). Despite significant advances in understanding size homeostasis, the influence of important physical parameters of the cell such as the turgor pressure and elastic properties of their outer envelope on size maintenance is not well known. Even though the molecules that control cell cycle and division have been identified (22), the ability to predict size from first principles remains a challenging problem.

Here, we develop an entirely different approach by casting the mechanism of size maintenance as an instability problem in hydrodynamics. We begin by studying the deformation response modes of the cell wall using a generalization of the Rayleigh-Plesset (RP) equation, which was derived in the context of modeling the dynamics of bubbles in fluids (23). The RP equation is a special case of the Navier-Stokes equation used to describe the size of a spherical bubble whose radius is *R*. We use the term “shell” generically, being equally applicable to membranes, and the composite layers making up the bacterial envelope or capsids. In our theory, the shell subject to deformation (e.g. expansion) exhibits two fundamental response modes: (*i*) elastic mode, where perturbative deformation of the cell wall is followed by initial size recovery, and (*ii*) unstable response where minute deformation results in continuous growth of the deformation. The initial size is not recovered in the unstable response mode, and hence we refer to it as the plastic response. This is similar to the yield point in springs beyond which original length of the spring is not recovered after stretching. The importance of these two fundamental deformation response modes in the context of bending and growth in rod shaped cells was investigated recently (24).

A key prediction of our theory is the relation between the deformation response modes and optimal size, dictated by a single dimensionless compliance parameter, *ζ*, expressed in terms of the elasticity of the shell and the turgor pressure. We show that an optimal cell size requires that *ζ* strike a balance between elastic and plastic response to deformation, thus maintaining microorganisms at the edge of stability. In general, from a biological perspective, remaining at the edge of stability might facilitate adaptation to changing environmental conditions. In fact, this principle is at the heart of ‘life at the edge’ with its consequences found from the molecular (25) to cellular level (26). The model consistently predicts the size of sphere-like bacteria and viruses given the physical properties of the cell and the protecting shell.

## II. THEORY

Approximating bacteria and viruses as bubbles with shells enables us to approach the problem of size maintenance using a generalized Rayleigh-Plesset (RP) equation (see Fig. 1). For a spherical bubble of radius, *R*(*t*), in a liquid at time *t*, the temperature and pressure outside the bubble, *T*_*out*_ and *p*_*out*_, are assumed to be constant. The liquid mass density, *ρ* and the kinematic viscosity, *ν*, are also taken to be constant and uniform. If we assume that the contents of the bubble are at a constant temperature and exert steady osmotic pressure on the bubble wall, *p*_*in*_, then the effect of the turgor pressure may be taken into account. We outline in Appendix A that the RP equation is motivated from the general equations governing fluid flow - the continuity and the Navier-Stokes equations. To extend the RP equation to study the size maintenance mechanism in microorganisms, an additional term for the bending pressure of the thin outer shell is required. The elastic energy (per unit area) of bending a thin shell is proportional to the square of the curvature (27). Thus, the generalized RP equation is,

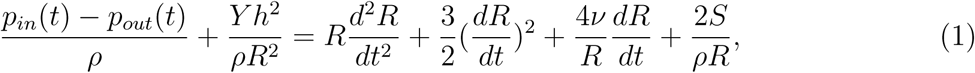

where *Y h*^2^*/R*^2^ is the bending pressure of the elastic shell, *Y* is the elastic modulus, *h* is the thickness of the shell, and *S* is the surface tension acting on the shell. The bending pressure, or the resistance to bending, arises due to the outer side of a bent material being stretched while the inner side is compressed (see Inset in Fig. 1). For more details on the bending pressure term see Appendix A. The first term on the left hand side accounts for the pressure difference between inside and outside of the cell and the other terms involve time derivatives of the radius.

**FIG. 1:**
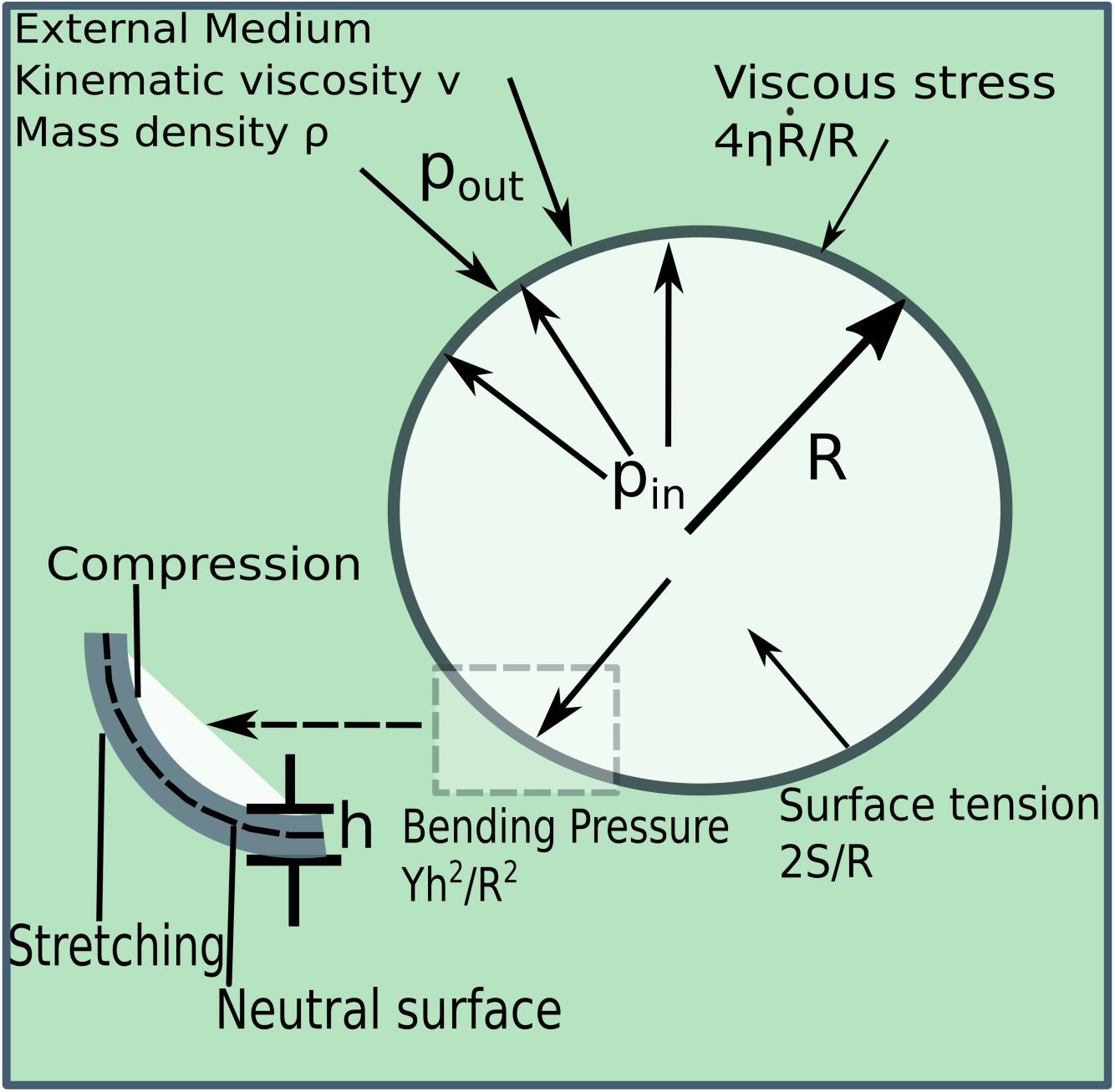
Illustration of the model cell based on the RP equation. The cell wall is a thin shell of thickness *h*. The stresses acting on the cell wall are labeled. Viscosity of the surrounding medium is *η* = *νρ*.

Shell displacement, *δR*(*t*), in the radial direction leads to *R*(*t*) = *R*_*e*_ + *δR*(*t*), where *R*_*e*_ is a constant. If *δR/R*_*e*_ ≪ 1, an equation for *δR*(*t*)*/R*_*e*_ may be derived,

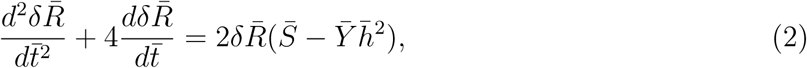

in non-dimensional units where 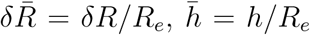 (see Appendix B for further details). The stretching energy per unit volume is, 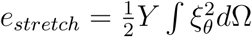, where *d*Ω is the differential solid angle, and the angular strain is defined as 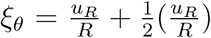 (28), *u*_*R*_ = *δR* is the displacement in the radial direction, giving rise to a second order contribution in terms of *δR* which we do not consider here. We choose 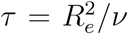 which sets the time unit and *R*_*e*_ (the mean cell size) is the unit of length. Similarly, the elastic modulus 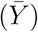 and surface tension are rescaled using 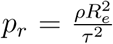 and 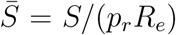. Three types of temporal behavior in 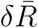 are illustrated in Fig. 2, where the radial displacement either increases, stays constant or decays. Both analytic and numerical solutions, with initial conditions 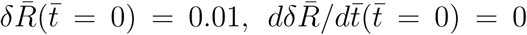, may be readily obtained, as detailed in Appendix C. Fig. 2a shows that as the dimensionless surface tension 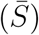 increases, the behavior of *δR*(*t*)*/R*_*e*_ changes from continuous decay to growth. In Fig. 2b, 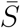 is kept constant while the stiffness of the shell is varied. Time dependent perturbative displacement, *δR*(*t*)*/R*_*e*_, once again shows three distinct trends as the shell stiffness, 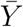, increases. Since *δR/R*_*e*_ is the strain experienced by the elastic shell due to infinitesimal deformation, these response modes signify a transition between the ‘elastic’ and the ‘plastic’ regime. The plastic regime corresponds to incremental growth in strain while in the elastic regime the strain decays to zero over time, implying that the cell size is maintained. The influence of the shell mechanical parameters on oscillation modes are discussed in Appendix D.

**FIG. 2:**
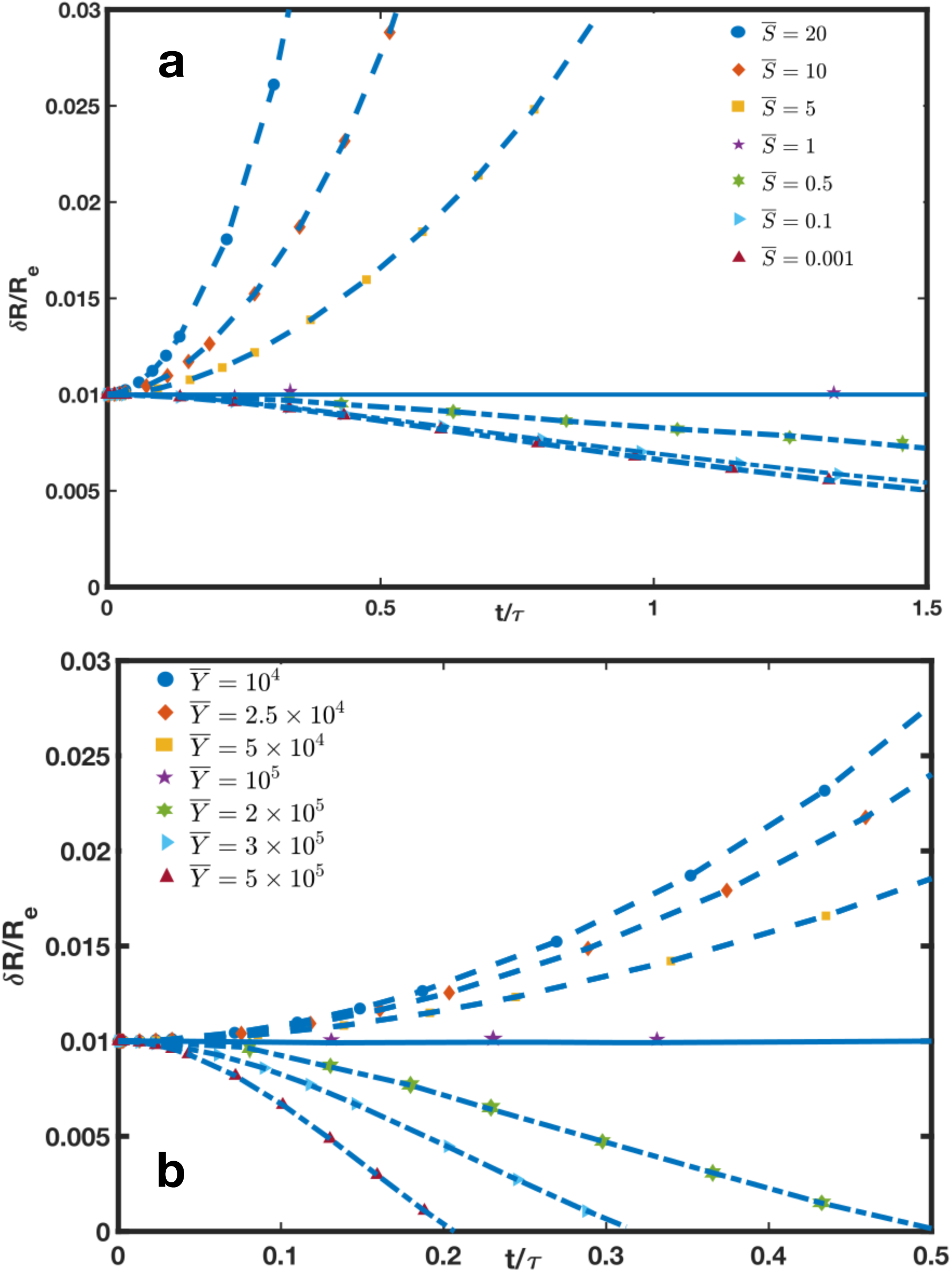
Solutions (a) for the first order RP equation shows the time dependent behavior of strain. Time is scaled by *τ* and length by *R*_*e*_. 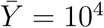 and 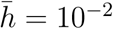 are kept constant while 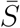 is varied. (b) Same as in (a) with 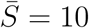 and 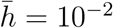 kept constant while 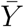 is varied.

The strain response modes depend on whether *S* is greater than or less than *Y h*^2^*/R*_*e*_ (see Eq. C2). If *S* > *Y h*^2^*/R*_*e*_, continuous growth in strain results in the ‘plastic’ regime. However, if *S* < *Y h*^2^*/R*_*e*_, a decaying solution for the strain leads to the ‘elastic’ regime. The critical value of the surface tension that dictates the boundary between the two regimes is predicted to be at 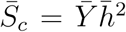. In Fig. 2a, for 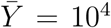 and 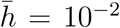, the critical surface tension corresponds to 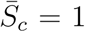. Similarly in Fig. 2b, we show that the critical elastic modulus is 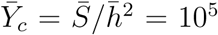, in agreement with numerical results. Note that Table I in Appendix E shows that the parameter ranges considered are physiologically relevant. Surface tension forces must be explicitly taken into account in studying envelope deformation of bacteria and viruses since the mechanical equilibrium of bacterial shells is determined by surface tension (29, 30). Similarly, mechanical properties of viral capsids are determined by surface tension (31) or effective surface tension-like terms (1).

**TABLE I:** Parameters used to obtain results in Fig 2 of the Main Text.

| Parameter | Value (Dimensionless) | Value (With Dimension) | Experimental Values |
| --- | --- | --- | --- |
| $R_e$ | — | $10^{-6}\text{m}$ | Fixed |
| $\nu$ | — | $1.0 \times 10^{-6} \text{m}^2/\text{s}$ (42, 43) | — |
| $\tau(R_e^2/\nu)$ | — | $1.0 \times 10^{-6} \text{sec}$ | — |
| $\rho$ | — | $10^3 \text{ kg/m}^3$ (42, 43) | — |
| $p_r(\frac{\rho R_e^2}{\tau^2})$ | — | $10^3 \text{ Pa}$ | — |
| $\bar{Y}(Y/p_r)$ | $10^4 - 5 \times 10^5$ | $10^7 - 5 \times 10^8 \text{ Pa}$ | See Table II |
| $\bar{S}(S/p_r R_e)$ | $10^{-1} - 20$ | $10^{-4} - 2 \times 10^{-2} \text{ N/m}$ | $10^{-2} - 10^{-1} \text{ N/m}$ (13, 29) |
| $\bar{h}(h/R_e)$ | $1.0 \times 10^{-2}$ | $1.0 \times 10^{-8}$ | See Table II |

An important prediction of the theory is that the dimensionless compliance parameter, *ζ*, quantifies the shell response to perturbative deformation and thereby sets a universal length scale for the size of microorganisms. The parameter *ζ* depends on intrinsic physical properties of the cell, which collectively play an important role in bacterial and viral shell deformation. The details of the derivation of,

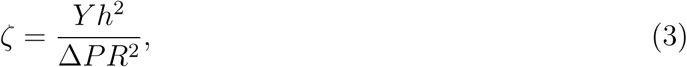

where Δ*P* = *p*_*in*_ − *p*_*out*_, are given in the Appendix B. The compliance parameter in Eq. 3 may be obtained by equating the bending pressure (∼ *Y h*^2^*/R*^2^ - this form is justified in the Appendix A) and the contribution arising from surface tension (∼ *S/R*, the last term in Eq. A9). By using the Young-Laplace equation for *S* ∼ Δ*PR*, we obtain Eq. 3.

Interestingly, the same parameter rewritten as, *κ* = 1*/ζ*, was found to be important in distinguishing between bending and tension-dominated response of the bacterial wall during the indentation of *Magnetospirillum gryphiswaldense* with an AFM tip (30). In a more recent study (24), the dimensionless variable *χ* (related to the compliance parameter by *χ*(*R/h*) = 1*/ζ*), was shown to demarcate the boundary between elastic and plastic bending regimes for cylindrical bacteria.

The elastic regime corresponds to *ζ* > 1, while for *ζ* < 1 the deformation is plastic. Since plastic and elastic deformation modes are expected to be of comparable importance in bacterial cell walls (24), we anticipate that the condition *ζ* = 1 could play an important role in determining size. Note that *ζ* = 1 corresponds to *δR*(*t*)*/R*_*e*_ = constant with neither decay nor growth in response to perturbative displacement. Thus, the boundary between the plastic and elastic regime (*ζ* = 1) lets us identify a critical radius,

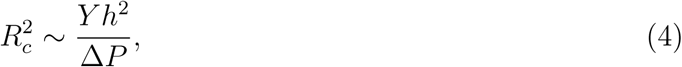

which is the central result of our work.

## III. ANALYSIS OF EXPERIMENTAL DATA

The predictions and the ensuing consequences of Eq. 4 are explored by analyzing experimental data. The critical radius obtained above unveils a universal dependence of the size of microorganisms on the intrinsic physical parameters of the cell and its outer shell – the pressure difference between inside and outside, and the elastic modulus and the thickness of the shell, respectively. We now analyze the size of bacteria (*S. aureus, E. coli, B. subtilis*), and viruses (*Murine Leukemia Virus (MLV)*, Φ*29 bacteriophage*, and *Human Immunodeficiency Virus (HIV)* etc) in relation to their shell physical properties. Data for the radius, elastic modulus, shell thickness and pressure difference were obtained from the literature. A comparison of the shell thickness, *h*, to the radial size, *R*, for 12 bacteria and viruses is presented in Fig. 3a. The thickness of the cell wall is,

**FIG. 3:**
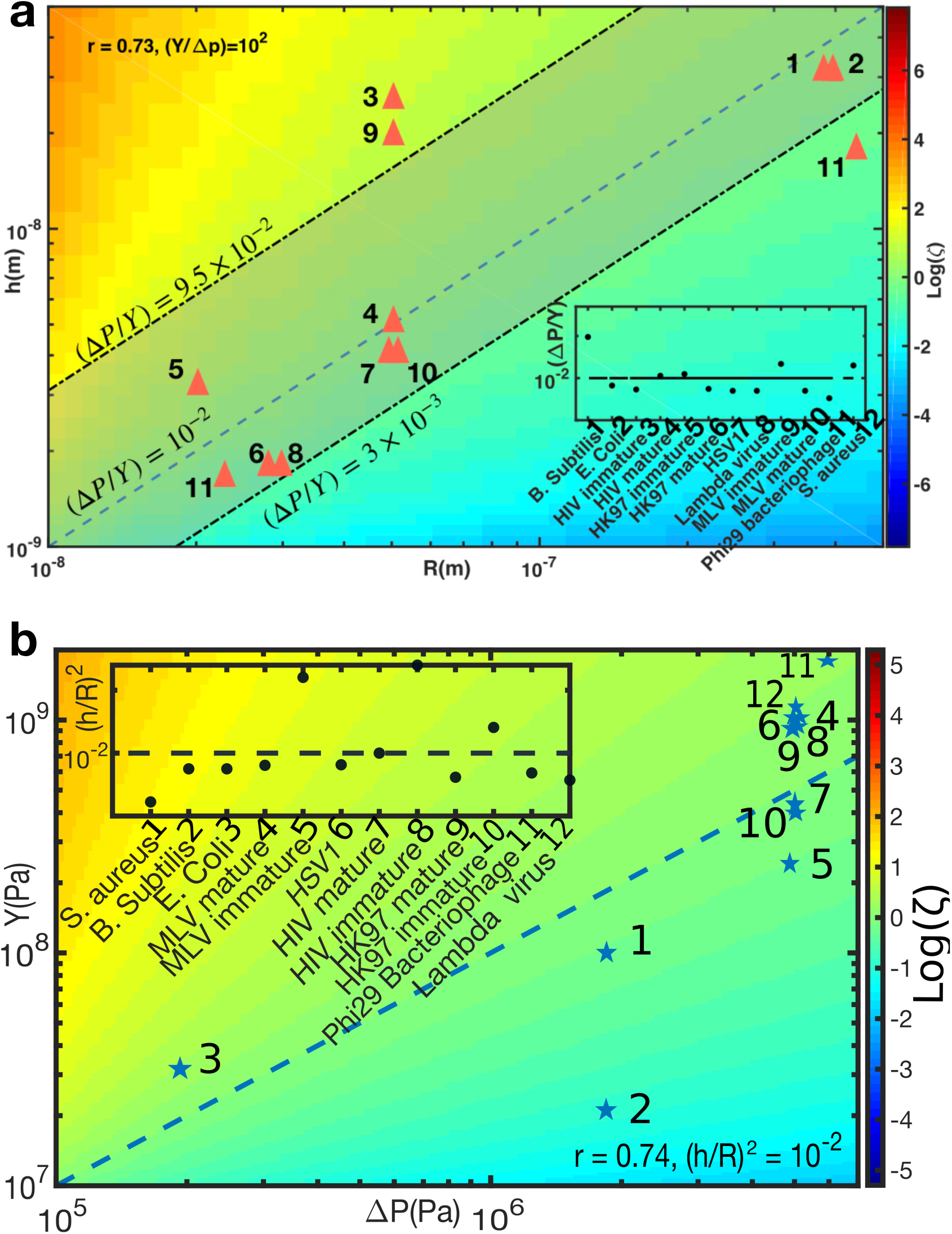
(a) Test of the relation given in Eq. (5) for different bacteria and viruses. Inset shows the ratio Δ*P/Y*. (b) Trend in shell elastic modulus (*Y*) versus turgor pressure (Δ*P*) in units of *Pa*. Inset shows the ratio of (*h/R*)^2^. The data were compiled from existing literature. Calculated *ζ* (in log scale, with fixed Δ*P/Y* = 10^−2^ in (a) and (*R/h*)^2^ = 10^2^ in (b)) is indicated in color heat map on the right of the figures. The dashed line in the figure marks *ζ* = 1.

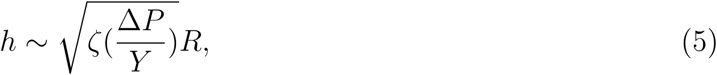

which is directly proportional to size (based on Eq. 3). Remarkably, the ratio of the turgor pressure to shell stiffness, Δ*P/Y* ∼ 10^−2^ (see Inset Fig. 3a), falls on a straight line for most of the bacteria and viruses. The ranges of Δ*P/Y* on the higher end from 9.5 × 10^−2^ for *B. subtilis* and on the lower end to 3 × 10^−3^, within an order of magnitude. The maximum observed value of Δ*P/Y* ∼ 9.5 × 10^−2^ and the minimum Δ*P/Y* ∼ 3 × 10^−3^ provides an upper and lower bound for our predicted *h* vs. *R* relationship, as plotted using dash-dot lines in Fig. 3a. In agreement with our theory, the shell thickness is linearly correlated to the overall size with the Pearson correlation coefficient, *r* = 0.73. As predicted by the theory, the data points lie close to *ζ* = 1 (marked by the dashed line) in Fig. 3a, indicative of the importance of the balance between plastic and elastic deformation modes in microorganisms.

Because correlation does not imply causation, we sought to test our prediction using an alternate set of variables. The relation between turgor pressure and shell elastic modulus can be predicted using,

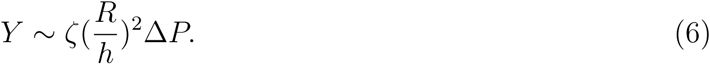

Taking the ratio between radial size and shell thickness, (*R/h*)^2^ ∼ 10^2^ (see Inset Fig. 3b), allows us to identify the preferable *ζ* regime. The Pearson correlation coefficient, *r* = 0.74, and the region where *ζ* = 1 is marked by a dashed line in Fig. 3b. When either Δ*P* or *Y* were not available, value of a similar species was used (see Table II in Appendix E for more details). Therefore, our conclusion that bacteria and viruses maintain their shell elastic properties to lie close to *ζ* = 1 is further borne out by this analysis.

**TABLE II:** Values of parameters used in the model. ∗For thickness of cell wall and membrane. Both in Tables I and II the numbers in brackets refer to reference citations.

| Organism | R (m) | h (m) | $\Delta P$(Pa) | Y(Pa) |
| --- | --- | --- | --- | --- |
| S. aureus | $4.4 \times 10^{-7}$(44) | $1.8 \times 10^{-8}$(44) | $1.9 \times 10^6$(44, 45) | $9.5 \times 10^7$(2) |
| B. subtilis | $4.0 \times 10^{-7}$(24) | $3.0 \times 10^{-8}$(24) | $1.9 \times 10^6$(24) | $2.0 \times 10^7$(24) |
| E. coli | $4.0 \times 10^{-7}$(24) | $3.0 \times 10^{-8}$ *(46) | $2.0 \times 10^5$(24) | $3.0 \times 10^7$(24) |
| Murine Leukemia Virus (mature) | $5.0 \times 10^{-8}$(33) | $4.0 \times 10^{-9}$(33) | $5.0 \times 10^6$(31) | $1.0 \times 10^9$(33) |
| Murine Leukemia Virus (Immature) | $5.0 \times 10^{-8}$(33) | $2.0 \times 10^{-8}$(33) | $5.0 \times 10^6$(31) | $2.3 \times 10^8$(33) |
| Herpes Simplex Virus 1 | $4.95 \times 10^{-8}$(1, 47) | $4.0 \times 10^{-9}$(1, 47) | $5.0 \times 10^6$(31) | $1.0 \times 10^9$(1, 47) |
| HIV (mature) | $5.0 \times 10^{-8}$(32) | $5.0 \times 10^{-9}$(32) | $5.0 \times 10^6$(31) | $4.4 \times 10^8$(32) |
| HIV (Immature) | $5.0 \times 10^{-8}$(32) | $2.5 \times 10^{-8}$(32) | $5.0 \times 10^6$(31) | $9.3 \times 10^8$(32) |
| HK97 (mature) | $2.8 \times 10^{-8}$(34) | $1.8 \times 10^{-9}$(34) | $5.0 \times 10^6$(31) | $9.0 \times 10^8$(34) |
| HK97 (Immature) | $2.0 \times 10^{-8}$(34) | $3.2 \times 10^{-9}$(34) | $5.0 \times 10^6$(31) | $4.0 \times 10^8$(34) |
| $\Phi 29$ bacteriophage | $2.3 \times 10^{-8}$(48) | $1.6 \times 10^{-9}$(48) | $6.0 \times 10^6$(48) | $1.8 \times 10^9$(48) |
| $\lambda$ virus | $2.95 \times 10^{-8}$(49) | $1.8 \times 10^{-9}$(1) | $5.0 \times 10^6$(31) | $1.0 \times 10^9$(49) |

### Maturation of Viruses

We propose that the physical mechanism of adapting to a specific value of *ζ* could be utilized by viral particles and bacteria to tightly maintain a specific size. We examine the consistency of this proposal by analyzing experimental data. We now explore the role of *ζ* (Eq. 3) in the viral maturation process. Double stranded (ds) DNA bacteriophages are known to undergo conformational and chemical changes that tend to strengthen the shell (1) by a process that resembles structural phase transition in crystals. This is necessary considering that the shells have to be able to withstand large internal pressures and at the same time be unstable so that their genome can be released into host cells during infection. The radius of a viral particle remains approximately constant throughout its life cycle. However, experimental evidence shows that shell thickness of viruses is tuned actively (see Table II in Appendix E and references therein). In *HIV, MLV* and *HK97* viruses, the shell thickness decreases during the maturation process (32–34). Interestingly, in *MLV* and *HK97* the decrease in shell thickness corresponds to an increase in capsid stiffness (33, 34). Given these clues, we quantify the role of *ζ* in the viral maturation process. Fig. 4 compares the size and shell elastic properties of individual viruses between their immature and mature phases. The tuning of the ratio of radius to the shell thickness to larger values as the virus matures is clearly observed (filled → hollow shapes). As the viral particle matures, a transition from the elastic regime(*ζ* > 1) to a plastic regime(*ζ* ≤ 1) is observed. The inset in Fig. 4 shows the fractional change in the compliance, *δζ/ζ* = (*ζ*_*immature*_ − *ζ*_*mature*_)*/ζ*_*immature*_. Notable change in *ζ* due to maturation occurs for all three viruses with *ζ*_*mature*_ ≪ *ζ*_*immature*_. This marks a crossover behavior in the viral lifecycle where surface tension-like forces in the shell begin to dominate the force associated with the shell stiffness. As before, the dashed line in the figure (*ζ* = 1) separates the elastic regime from the plastic regime. We surmise that such an adaptation is necessary for the onset of instability in viral capsids.

**FIG. 4:**
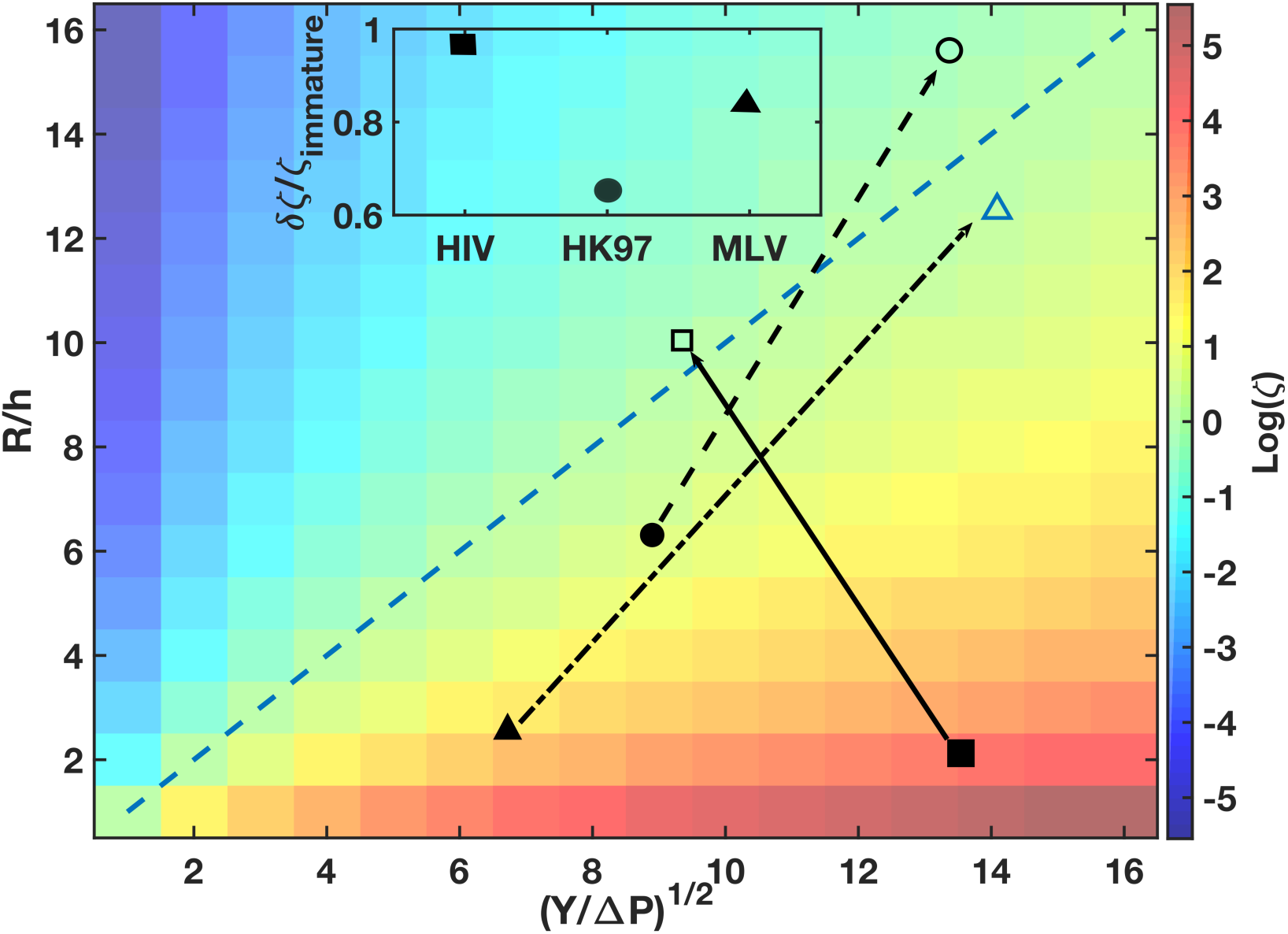
Comparing the ratio of *R/h* to (*Y/*Δ*P*)^1/2^ during the immature and mature stages of viruses. Mature/Immature HIV (□) (32), Mature/Immature HK97 virus (○) (34, 35), Mature/Immature Murine Leukemia Virus (MLV △) (33). Hollow and filled symbols correspond to mature and immature viruses respectively (with lines joining them as guides to the eye). Calculated *ζ* is indicated in color scale on the right. The dashed diagonal line in the figure marks *ζ* = 1. Lines connecting filled to hollow symbols visualize the tuning of the physical parameters as a virus matures.

The elastic modulus (*Y*) of the viral shell is an important parameter in the maturation process of viruses. Shells of *MLV* and *HK97* become stiffer as they mature. As a result of the increased shell stiffness, *h* decreases while maintaining approximately the same size. In the three different viruses analyzed, *ζ* approaches 1, which we propose to be a general property of viral maturation.

### Bacteria

We explore further the consequence of adapting to a specific value of *ζ* in a bacterium. The thickness of the *S. aureus* cell wall is tuned to higher values as a result of nutrient depletion in the stationary phase (in *S. aureus* synthetic medium) (36). Glycine depletion in the nutrient medium forces *S. aureus* to make “imperfect” peptidoglycan resulting in a less rigid cell wall, which is more susceptible to lysis (37). *S. aureus* responds by increasing the peptidoglycan thickness. The observed adaptation behavior of the cell wall thickness is to be expected from Eq. (4). Given a decrease in *Y* (due to a defective cell wall) and assuming that the ratio of pressure difference to *Y* is a constant, the bacteria can maintain its size by increasing the thickness, *h*. We note that in the stationary phase bacteria arrest their growth and enter dormancy (38), maintaining a constant size *R*.

A scaling behavior obtained earlier by balancing the bending pressure of shells with turgor pressure (12) showed that *R/h* is proportional to (*Y/*Δ*P*)^1/3^, focusing on alga cells and fungi. This proposed scaling arises due to surface tension-like forces not being considered, which perhaps accounts for the departure from the proposed scaling for bacteria and viruses (the focus of this study). We quantitatively compare the best fit to experimental data presented in Fig. 3a to the two different scaling behaviors: (i) 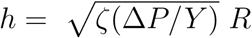, and (ii) *h* = *α*^−1^(Δ*P/Y*)^1/3^ *R* (12) and show that the experimental data is better accounted by the generalized RP theory (see Appendix F).

## IV. CONCLUSION

By generalizing a hydrodynamic model based on the Rayeigh-Plesset equation, originally formulated in the context of bubbles in fluids, we proposed a novel unified framework to predict the size of bacteria and viruses from first principles. Given the shell elastic properties and the pressure differential between the inside and outside, the importance of selecting a deformation response mode is shown as a possible mechanism to constrain size. Nanoscale vibrations, proposed as a signature of life (39), could provide a natural basis for bacteria and viruses to detect the elasticity of shells. We identified a compliance parameter, *ζ*, in terms of the physical properties of the cell as the most relevant variable controlling cell size. Viral particles are especially sensitive to *ζ*, and we predict that shell properties evolve to minimize *ζ* during maturation. By merging approaches from hydrodynamics and elasticity theory, we have proposed a new mechanism for an important question in cell biology pertaining to size regulation. In conjunction with studies on the role of biochemical processes in shape and size maintenance, the importance of physical parameters should also be considered in order to fully understand size homeostasis in bacteria and viruses.

## V. APPENDIX

### Appendix A: Motivation of Rayleigh-Plesset Equation

We begin with the equations governing fluid flow in the presence of a bubble (Fig. 1 in the Main Text), the continuity equation,

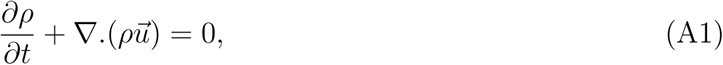

and the Navier-Stokes equation,

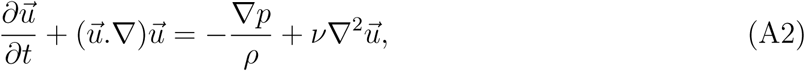

where 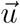 is the velocity field. Other parameters are defined in the main text. Considering the incompressible fluid limit, the continuity equation requires that,

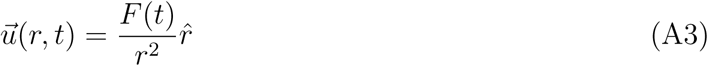

where *F* (*t*) can be related to *R*(*t*) by a boundary condition at the bubble surface. Under the assumption that there is no mass transport across the interface of the bubble, *u*(*R, t*) = *dR/dt*, and we obtain 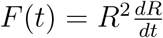. Since we are interested only in the radial direction, for simplicity, we drop the vector sign henceforth. Substituting *u* from Eq. (A3) into the the Navier-Stokes equation above gives,

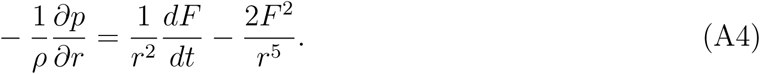

To proceed further, we consider a small, thin segment of the bubble liquid interface and the net forces (per unit area) acting along the radial direction. They are:

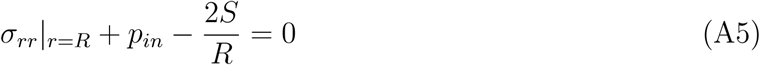

where *σ*_*rr*_ is the stress tensor in the fluid, and *S* is the surface tension of the bubble film. For a Newtonian fluid, 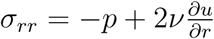, and we obtain

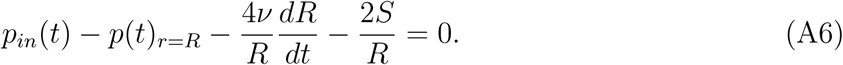

Integrating Eq. (A4) gives the pressure at the bubble surface and following the same steps as in chapter 4 of Ref. (23), the generalized Rayleigh-Plesset (RP) equation is,.

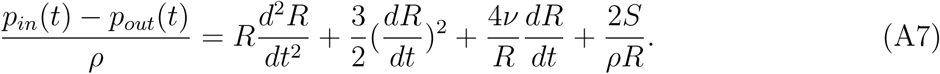

This equation, in the absence of surface tension and viscous terms, was first derived by Lord Rayleigh (40) in 1917, and was first applied to the study of cavitation bubbles by Plesset (41).

In order to generalize Eq. (A7) to investigate size maintenance mechanism in bacteria, an additional term accounting for the bending pressure of the thin outer shell is needed. The elastic energy (per unit area) of bending is proportional to the square of the curvature of the thin shell (12). The bending pressure, *p*_*b*_, of the elastic shell is

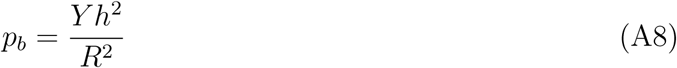

where *Y* is the elastic modulus, and *h* is the shell thickness. Therefore, the generalized RP equation that we shall utilize in this study is:

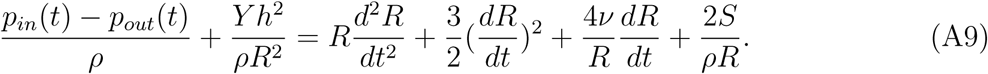

Stresses acting on the shell are visualized in Fig. 1 of the Main Text.

#### Bending pressure term

Here, we motivate why the bending pressure term should scale as *Y h*^2^*/R*^2^. Consider a thin sheet of material with sides of length *L*, thickness *h* and bend it so that it develops a radius of curvature, *R*. As illustrated in Fig. 1 of the Main Text, one side of the material will be stretched while another side is compressed in order to accommodate the bending. Assuming that the bending energy of the thin sheet is given by,

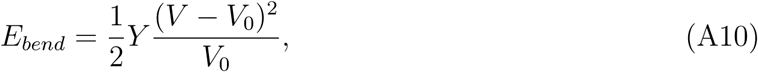

where *Y* is the elastic modulus of uniaxial extension or compression, *V* is extended or compressed volume and *V*_0_ the initial volume. Then, the bending energy per unit volume is

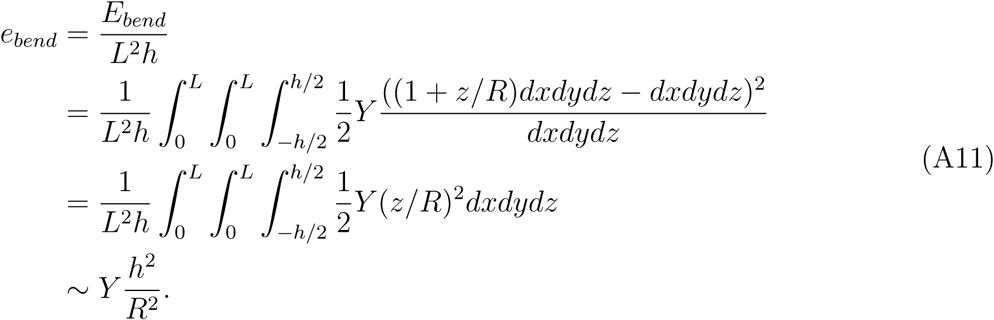

Here, the extended or compressed volume element is

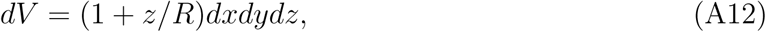

for a thin sheet bent along the z-direction with radius of curvature, *R*. Note that one of the integrals is over the thickness of the shell, *h*.

Moreover, Eq. (2) of Ref. (12) proposes that elastic energy **per unit area** for bending is,

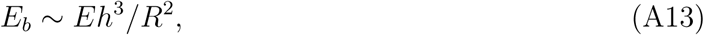

where *E* = *Y*, the elastic modulus. To obtain the corresponding bending pressure, with units of energy per unit volume,

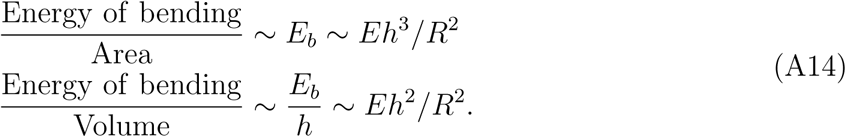

The relevant volume is that of the shell, given by *Area* × *h*. The key point to note here is that the volume contribution comes from the integral over the thickness, *h*, of the shell.

### Appendix B: Perturbation Theory

Having motivated the origin of the generalized RP equation, we now investigate the time dependent behavior of radial displacement. We consider the displacement of the spherical shell along the radial direction as a perturbation,

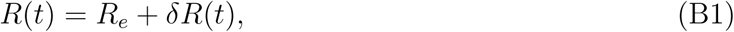

with *R*_*e*_ being a constant and *δR/R*_*e*_ ≪ 1. By substituting Eq. (B1) into Eq. (A9) we obtain,

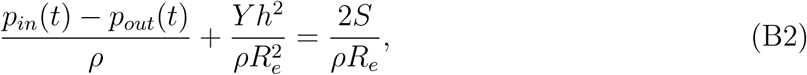

the generalized RP equation to zeroth order. To first order in *δR* the generalized RP equation takes the form:

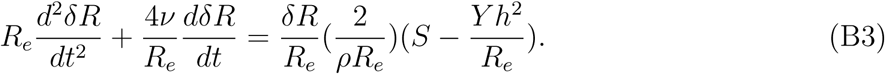

Note that the right hand side of Eq. (B3) has the same sign as the perturbation (*δR* > 0) if

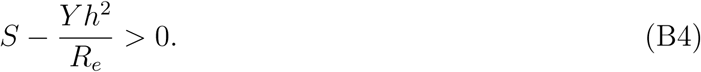

If the condition above holds, the velocity and acceleration of radial growth have the same sign as the perturbation implying that any small deviation in the radius will cause *R*(*t*) to deviate farther away from *R*_*e*_. We refer to this as the plastic regime. Elastic regime is obtained, to linear order, if the opposite condition, 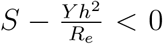 holds. Using the zeroth order Eq. B2, we obtain

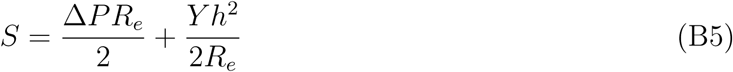

where Δ*P* = *p*_*in*_ − *p*_*out*_ is the differential pressure between inside and outside. Substituting *S* into Eq. B4 leads to the crucial compliance scale,

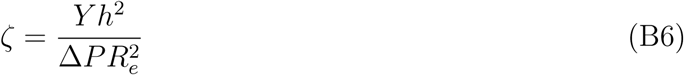

such that *ζ* < 1 corresponds to *S* − *Y h*^2^*/R*_*e*_ > 0 resulting in the plastic regime, and *ζ* > 1 leads to the elastic regime.

The critical radius *R*_*c*_ = *Y h*^2^*/S* that separates the plastic and elastic regime can now be re-written as,

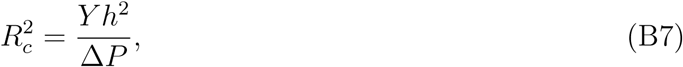

which is equivalent to *ζ* = 1.

### Appendix C: Solutions

Prior to presenting the details of the solution to the RP equation, we present a summary of the units. The characteristic time scale, *τ*, is defined in terms of the kinematic viscosity as 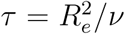, where *R*_*e*_ rescales the length *R*. All pressure terms and surface tension are rescaled using 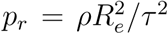 and 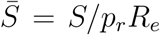 respectively. Using 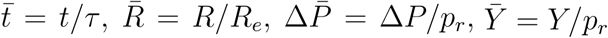, and 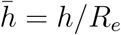 one obtains from Eq. (A9) the dimensionless RP equation,

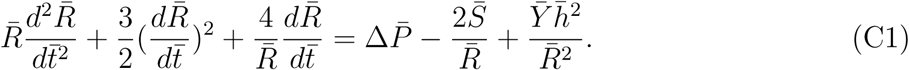

The first order RP equation (Eq. (B3)) in the dimensionless form is,

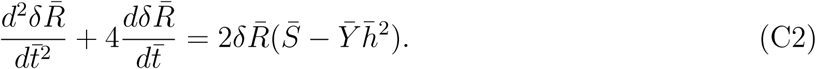

In order to solve the equations above, the values of the parameters such as 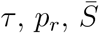 are needed. The arbitrary length scale is chosen to be *R*_*e*_ ∼ 10^−6^m as we focus on the size of microorganisms. We use the kinematic viscosity of water as a typical value for *ν* ∼ 10^−6^m^2^/s (42, 43), and for mass density *ρ* ∼ 10^3^kg/m^3^. For the surface tension per unit length, we use *S* ∼ 19nN*/μ*m (13). The magnitude of the quantities *τ* ∼ 10^−6^s, *p*_*r*_ ∼ 10^3^N/m^2^ are thus obtained. The initial conditions are 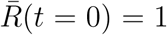 and 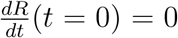. We use the typical values of 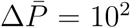 and 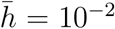 for bacteria (see Table I for further details). The numerical solution (using MATLAB) for 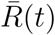 from Eq. C1 above is presented in Fig. 5. The first order Eq. (C2) is solved both analytically and numerically using MATLAB (ode15s solver). The analytical solution to first order RP equation is,

**FIG. 5:**
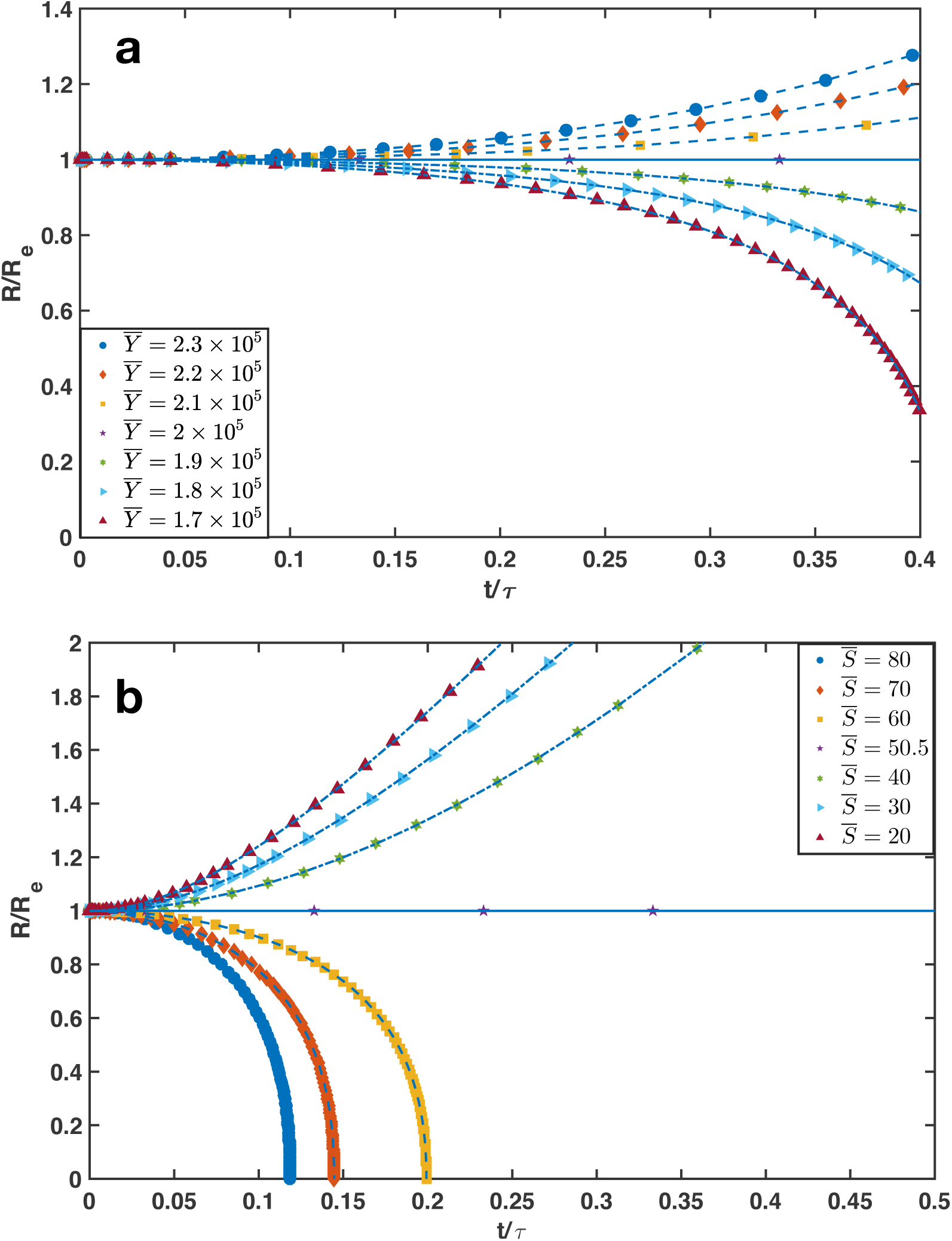
Numerical solutions: (a) Time dependence of the total size 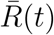 from the dimensionless RP equation. Time is scaled by the parameter *τ* and length by *R*_*e*_. 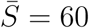 and 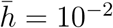 are kept constant while 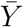 is varied. (b) As for (a) but 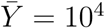 and 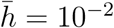 are constant while 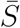 is changed.

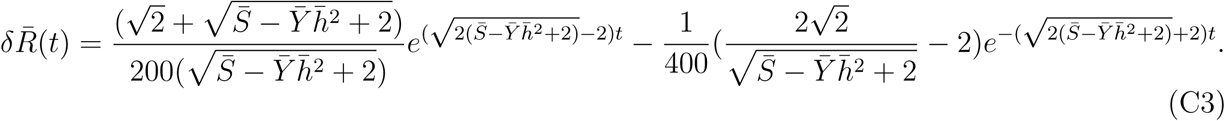

For 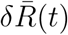, the radial displacement strain, two behaviors are observed: growth as a function of time corresponding to the plastic regime and an elastic regime where 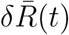 decays as a function of time. A similar behavior is also seen for 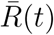.

The bending pressure drives the growth in 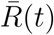 with larger 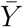 leading to continuous increase in size. Change in the behavior of 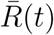 from continuous growth to decay either due to a decrease in elastic modulus or increase in surface tension of the cell wall is observed. This can be understood from the sign of the right hand side of Eq. C1, validating the zeroth order Eq. B2.

### Appendix D: Oscillation Modes

To study the oscillation modes of the bacterial and viral shell, we substitute *δR* = *δR*_*a*_*e*^*iωt*^ (*ω* is the oscillation frequency) into Eq. (B3) and obtain

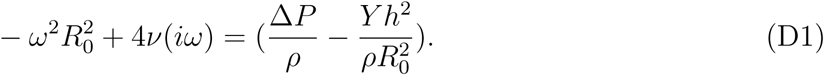

Solving for the frequency of the normal modes,

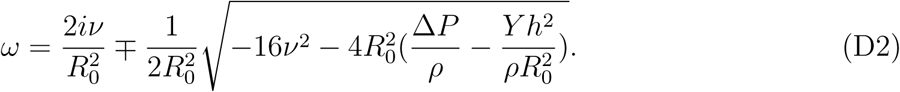

The real part of *ω* (*ω*_*R*_) must exist for an oscillatory resonant mode to be present. Focusing on the low viscosity limit (*ν* → 0), the resonant oscillation modes can only exist in the elastic regime with *ζ* > 1, which is illustrated in Fig. 6. Defining 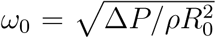, the dimensionless oscillation frequency of the shell is.

**FIG. 6:**
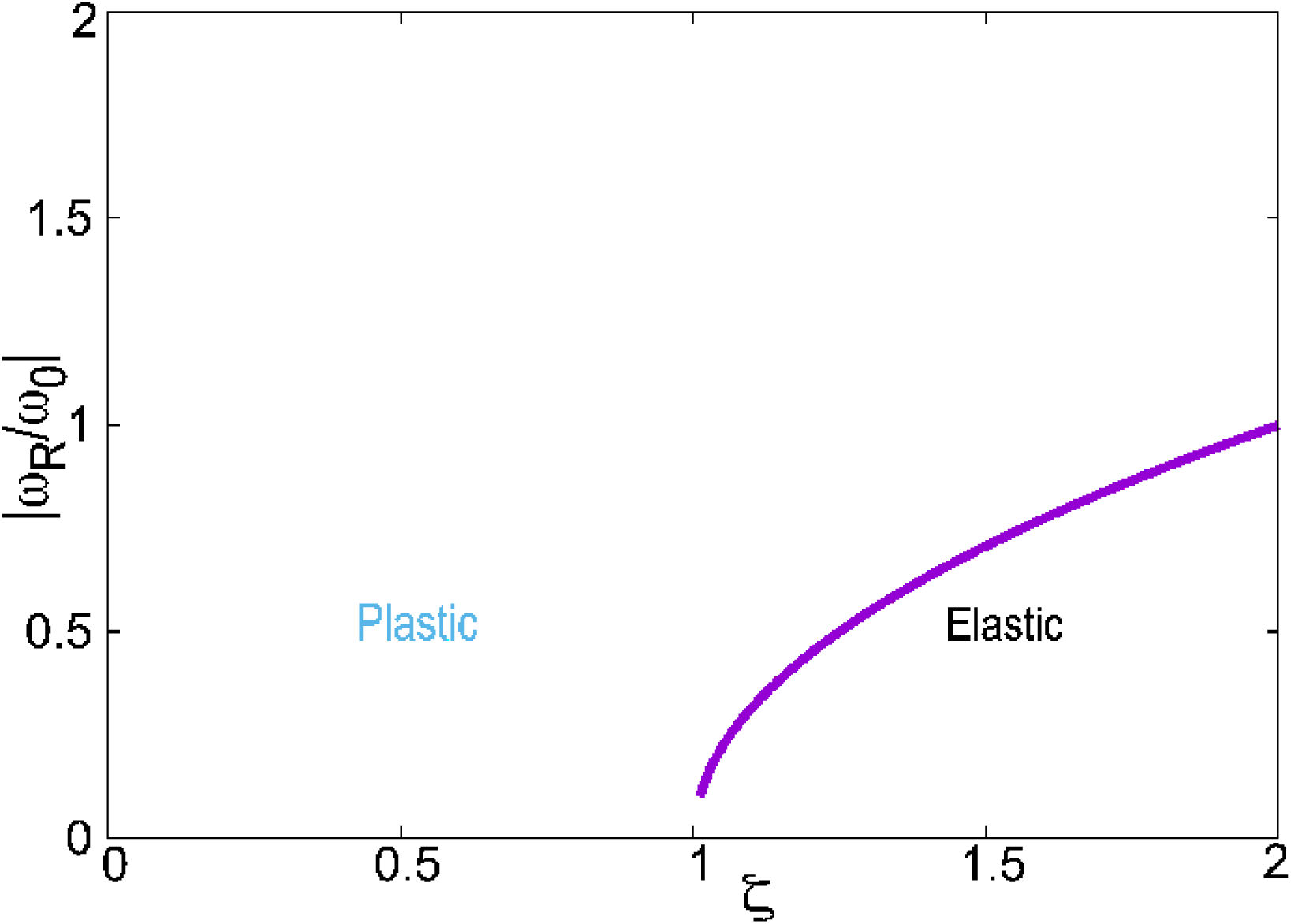
Plot showing the relation between the resonant oscillatory mode and *ζ*. The boundary between the elastic and plastic regimes is indicated. Undamped vibrational shell response is not expected to exist in the plastic regime.

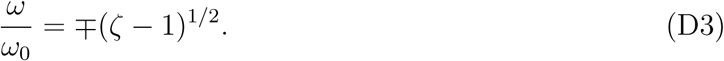

### Appendix E: Data

Numerical values for the various physical parameters considered in Fig 2 of the Main text are summarized in Table I. The values considered for the length and time scale, elastic modulus, surface tension are all shown to be in the relevant physiological range for bacteria.

We summarize the data in Table II for the various physical parameters characterizing bacteria and viruses used to illustrate our theory.

### Appendix F: Comparison to An Existing Model

By analyzing the data in Fig. 3 (Main Text), we assess whether the variability/spread in data allows us to compare our theory and the one proposed by Boudaoud (12). Two different scaling laws are put to test: (i) 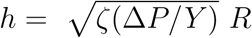, and (ii) *h* = *α*^−1^(Δ*P/Y*)^1/3^ *R* (*α* being a constant (12)). Even though the variation of *h* with (Δ*P/Y*) is very different between the two models, the scaling laws relating thickness of the shell, *h*, and cell size, *R*, are of the form,

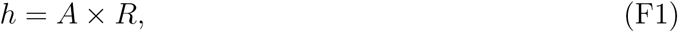

where the coefficient *A* (*A*_*B*_ = *α*^−1^(Δ*P/Y*)^1/3^ or 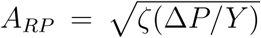) is to be determined. Fitting the data based on the theory proposed in Ref. (12) was found to be quantitatively worse (see Fig. 7), based on residuals, 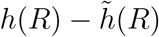 and percentage errors. Here, 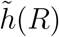 represents the shell thickness values predicted from theory.

**FIG. 7:**
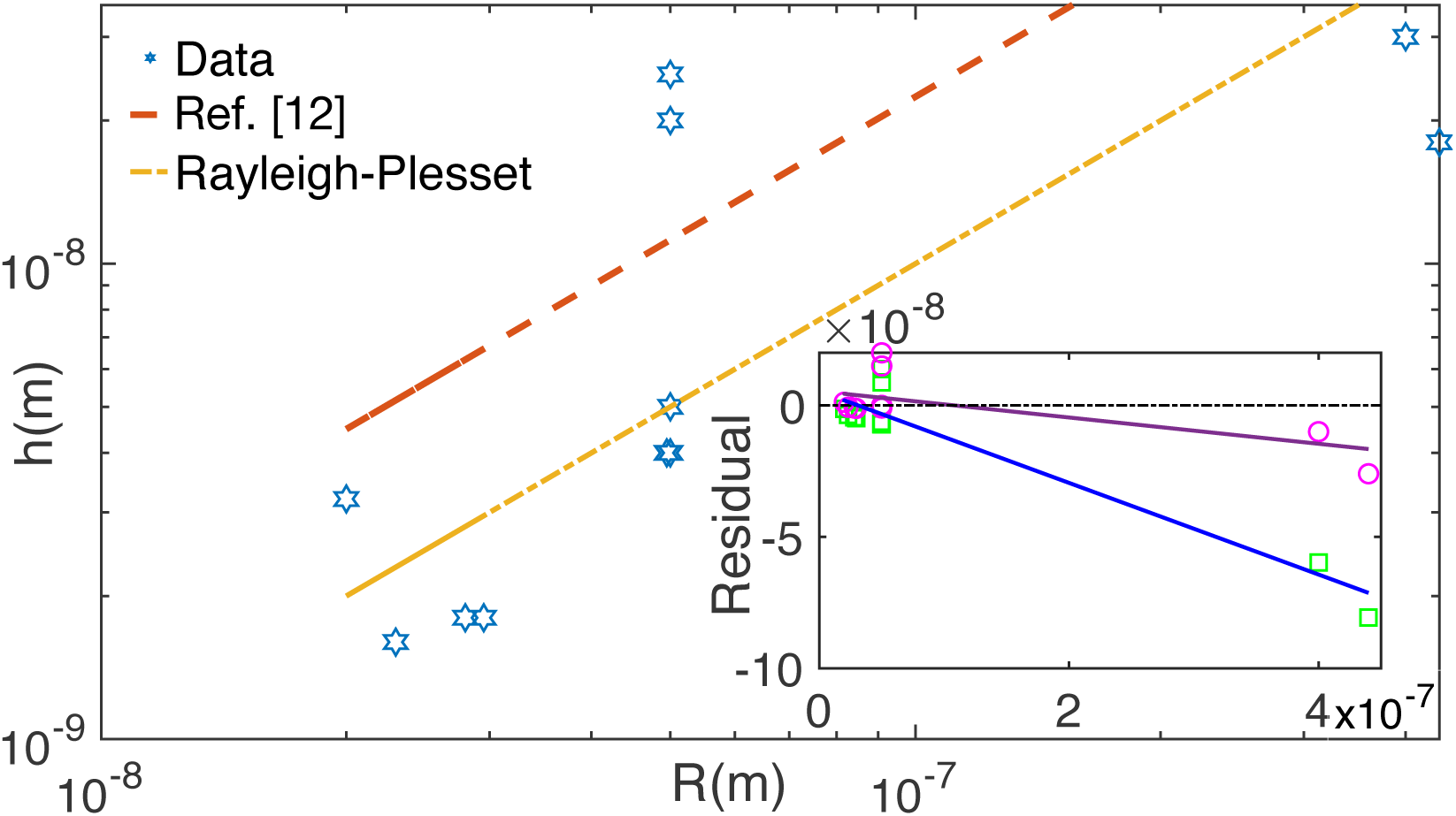
Shell thickness (*h*) for organisms of radial size *R*. The red dashed line is the proposed scaling of shell thickness according to the theory proposed by Ref. (12). The orange dashed-dot line is calculated from the generalized Rayleigh-Plesset theory. Inset: Residuals from theory in Ref. (12) (square) and generalized Rayleigh-Plesset theory fit (circle) as a function of *R*. Residuals are fit merely to function as guides for the eye.

The coefficient *A* is a function of the parameter Δ*P/Y*. As shown in the inset of Fig. 3a (Main Text), for the microorganisms considered, the ratio of the turgor pressure to shell stiffness is well approximated by Δ*P/Y* ∼ 10^−2^. Therefore, we estimate the coefficient *A* and compare it to best fit of the experimental data. For the experimental data, best fit to shell thickness versus *R* is obtained with coefficient *A* = 0.053 with a 95% confidence interval (CI) spanning the lower limit of 0.0198 and an upper limit of 0.0858.

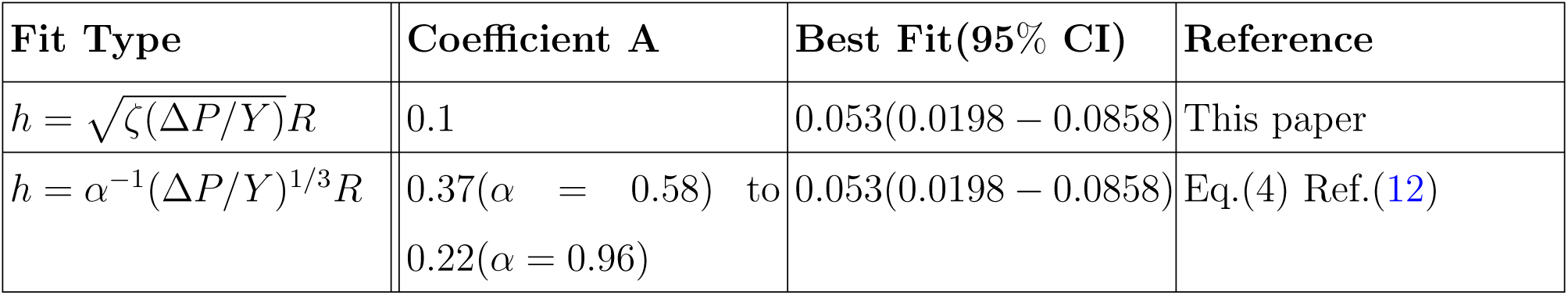

Error analysis reveals that the scaling relation proposed by our generalized Rayleigh-Plesset model gives a better agreement with experimental data, which is likely due to the importance of the surface tension. Considering the value of *A* = 0.0858 at the upper limit of the CI, we estimate the error Δ (expressed as a %) as,

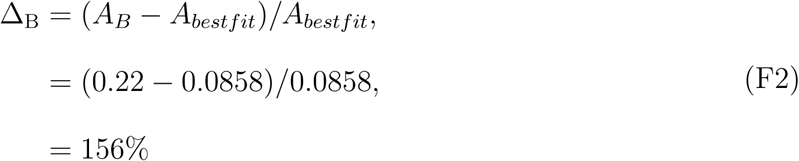

For the generalized R-P theory, the error is given by,

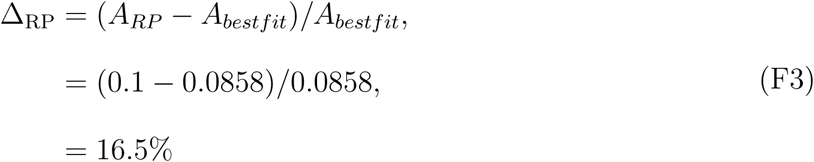

Therefore, quantitatively we conclude that the experimental data for the sizes of bacteria and viruses are better accounted for by the generalized Rayleigh-Plesset theory proposed here. For microscopic cell sizes of *R* < 1*μm*, Ref. (12) notes that the experimental data departs from the theoretical scaling while good agreement is observed for cells of size in the range 1 − 100*μm*. Finally, it will be most interesting to perform experiments to measure *h* by changing Δ*P* while keeping *Y* constant or vice versa. This would distinguish between the generalized RP prediction and the 1/3 scaling proposed in Ref. (12).

## Acknowledgements

This work was supported by the National Science Foundation through NSF Grant No. PHY 17-08128. Additional support was provided by the Welch Foundation through the Collie-Welch Chair (F-0019). We are grateful to Mauro Mugnai, and Xin Li for discussions and comments on the manuscript.

## Data availability

The authors declare that the data supporting the findings of this study are available within the paper [and its supplementary information files].

